# A Specialized Centrosome-Proteasome Axis Mediates Proteostasis and Influences Cardiac Stress through Txlnb

**DOI:** 10.1101/2024.02.12.580020

**Authors:** Jared M. McLendon, Xiaoming Zhang, Colleen S. Stein, Leslie M. Baehr, Sue C. Bodine, Ryan L. Boudreau

## Abstract

**Background:** Centrosomes localize to perinuclear foci where they serve multifunctional roles, arranging the microtubule organizing center (MTOC) and anchoring ubiquitin-proteasome system (UPS) machinery. In mature cardiomyocytes, centrosomal proteins redistribute into a specialized perinuclear cage-like structure, and a potential centrosome-UPS interface has not been studied. Taxilin-beta (Txlnb), a cardiomyocyte-enriched protein, belongs to a family of centrosome adapter proteins implicated in protein quality control. We hypothesize that Txlnb plays a key role in centrosomal-proteasomal crosstalk in cardiomyocytes.

**Methods:** Integrative bioinformatics assessed centrosomal gene dysregulation in failing hearts. Txlnb gain/loss-of-function studies were conducted in cultured cardiomyocytes and mice. Txlnb’s role in cardiac proteotoxicity and hypertrophy was examined using CryAB-R120G mice and transverse aortic constriction (TAC), respectively. Molecular modeling investigated Txlnb structure/function.

**Results:** Human failing hearts show consistent dysregulation of many centrosome-associated genes, alongside UPS-related genes. Txlnb emerged as a candidate regulator of cardiomyocyte proteostasis that localizes to the perinuclear centrosomal compartment. Txlnb’s interactome strongly supports its involvement in cytoskeletal, microtubule, and UPS processes, particularly centrosome-related functions. Overexpressing Txlnb in cardiomyocytes reduced ubiquitinated protein accumulation and enhanced proteasome activity during hypertrophy. Txlnb-knockout (KO) mouse hearts exhibit proteasomal insufficiency and altered cardiac growth, evidenced by ubiquitinated protein accumulation, decreased 26Sβ5 proteasome activity, and lower mass with age. In Cryab-R120G mice, Txlnb loss worsened heart failure, causing lower ejection fractions. After TAC, Txlnb-KO mice also showed reduced ejection fraction, increased heart mass, and elevated ubiquitinated protein accumulation. Investigations into the molecular mechanisms revealed that Txlnb-KO did not affect proteasomal subunit expression but led to the upregulation of Txlna and several centrosomal proteins (Cep63, Ofd1, and Tubg) suggesting altered centrosomal dynamics. Structural predictions support Txlnb’s role as a specialized centrosomal-adapter protein bridging centrosomes with proteasomes, confirmed by microtubule-dependent perinuclear localization.

**Conclusions:** Together, these data provide initial evidence connecting Txlnb to cardiac proteostasis, hinting at the potential importance of functional bridging between specialized centrosomes and UPS in cardiomyocytes.

## 1. Introduction

Centrosomes are membrane-less cell organelles composed of centrioles and their satellite proteins, that comprise the microtubule organizing center (MTOC) and perform a variety of cellular processes, the most studied being cell division. An oft-overlooked feature of the centrosome is its association with proteasomes^1^. Proteasomes and ubiquitin are enriched at centrosomes, proteasomes fraction in sucrose gradients with centrosome proteins, and centrosomes double in size when proteasomes are inhibited. In addition, misfolded proteins can accumulate at the centrosome even when proteasomes are functional^2^. Together, these data suggest that centrosomes are a central reservoir where misfolded and aggregated proteins traffic to and accumulate, leading to efficient degradation and removal of unwanted proteins^3^.

Terminally differentiated cardiomyocytes are a unique cell type with specialized cell biology to enable rhythmic, synchronous contraction sufficient to maintain cardiac output. Adult cardiomyocytes lack centrosomes and instead undergo centrosome reduction, through which centriolar and satellite proteins redistribute from discrete foci into cage-like structures surrounding nuclei^4^ via incompletely understood mechanisms. Along with this specialized centrosome, the MTOC and a large population of proteasomes also relocate to this perinuclear space. Whereas proteasomal insufficiency contributes to several types of heart failure, the specific contribution of centrosome-associated proteasomal pools has not been explored. In addition, the specific pathways and players that regulate centrosome reduction in cardiomyocytes are incompletely understood but involve AKAP6/9 dependent recruitment and nuclear anchoring of pericentriolar proteins,^5,6^ loss of CEP135 proteins, and alternative PCNT splicing.^7^ Beyond this, failure of cardiomyocytes to undergo centrosome reduction leads to immature cardiomyocyte phenotypes with disorganized cytoskeleton and impaired contractility.^8,9^ This suggests that specialized centrosomes play critical roles in maintaining cardiac homeostasis, yet the overall mechanisms and links to heart failure remain understudied.

In this study, we found broad dysregulation of genes encoding centrosomal proteins in human heart disease. We stumbled upon the taxilin gene/protein family (Txlna, Txlnb, and Txlng)^10^ that encodes centrosomal adapter proteins.^11,12^ One member, Txlnb, is highly enriched in cardiomyocytes, but its roles in cardiac biology and disease remain understudied^13–15^. Notably, we found that Txlnb influences protein homeostasis in cardiomyocytes. Txlnb overexpression promotes increased proteolysis through the ubiquitin-proteasome system (UPS) and increased protein synthesis, perhaps to increase turnover and replacement of cardiac proteins. In contrast, genetic knockout of Txlnb (Txlnb-KO) in mice causes proteasomal insufficiency, coincident with accumulation of ubiquitinated protein conjugates and reduced proteasome activity. Txlnb-KO mice exhibit stunted cardiac growth with age, exacerbated proteotoxicity-induced heart failure, and worse responses to pressure-overload along with dysregulated centrosomal protein expressions. Overall, our study sheds light on the intricate interplay between centrosomes, the UPS, and Txlnb in maintaining cardiac homeostasis, highlighting an understudied centrosome-proteasome interface that can influence cardiac stress.

## 2. Results

### 2.1 Centrosome-associated genes are broadly dysregulated in heart failure

To explore new and understudied genes/pathways involved in the cellular pathology underlying human heart failure, we performed a meta-analysis using publicly available RNA sequencing data **(Supplement Table 1)**. We identified 2,885 genes that are significantly downregulated (beta < −0.2, adj. pval < 0.05) in at least three of four independent datasets, and subsequent gene ontology analysis of this gene set (using Enrichr web server^16^) revealed functional enrichments related to UPS, centrosomes, and MTOC **(Figure 1a, Supplement Table 2)**. We next intersected a list of centrosome-associated genes, defined by gene ontology annotations (microtubule organizing center (GO:0005815), the centrosome (GO:0005813), and centriole (GO:0005814)) and found 43 upregulated and 128 downregulated genes **(subset shown in Figure 1b, Supplement Table 3)**, suggesting bi-directional rewiring of the centrosome in heart failure. To explore this centrosome association beyond ontology-curated lists, we next examined public data from centrosome-targeted biotin proximity labeling studies^17–21^ and identified the taxilin protein family among several other understudied genes. From one experiment with 22 different centrosome baits^17^, we noted that Txlna and Txlng show significant enrichment with ∼50% of baits tested, whereas Txlnb showed ∼25-fold enrichment to one centrosome bait (Cep63) with trending significance (fdr=0.14) **(Supplement Figure 1a-c)**. Notably, this minimal Txlnb enrichment across the baits likely relates to its low expression level (TPM=0.4) in HEK293-derived cells (used in bait screening), as Txlnb expression is known to be highly enriched in skeletal and cardiac myocytes^13^ **(Supplement Figure 1d)**.

**Figure 1.**
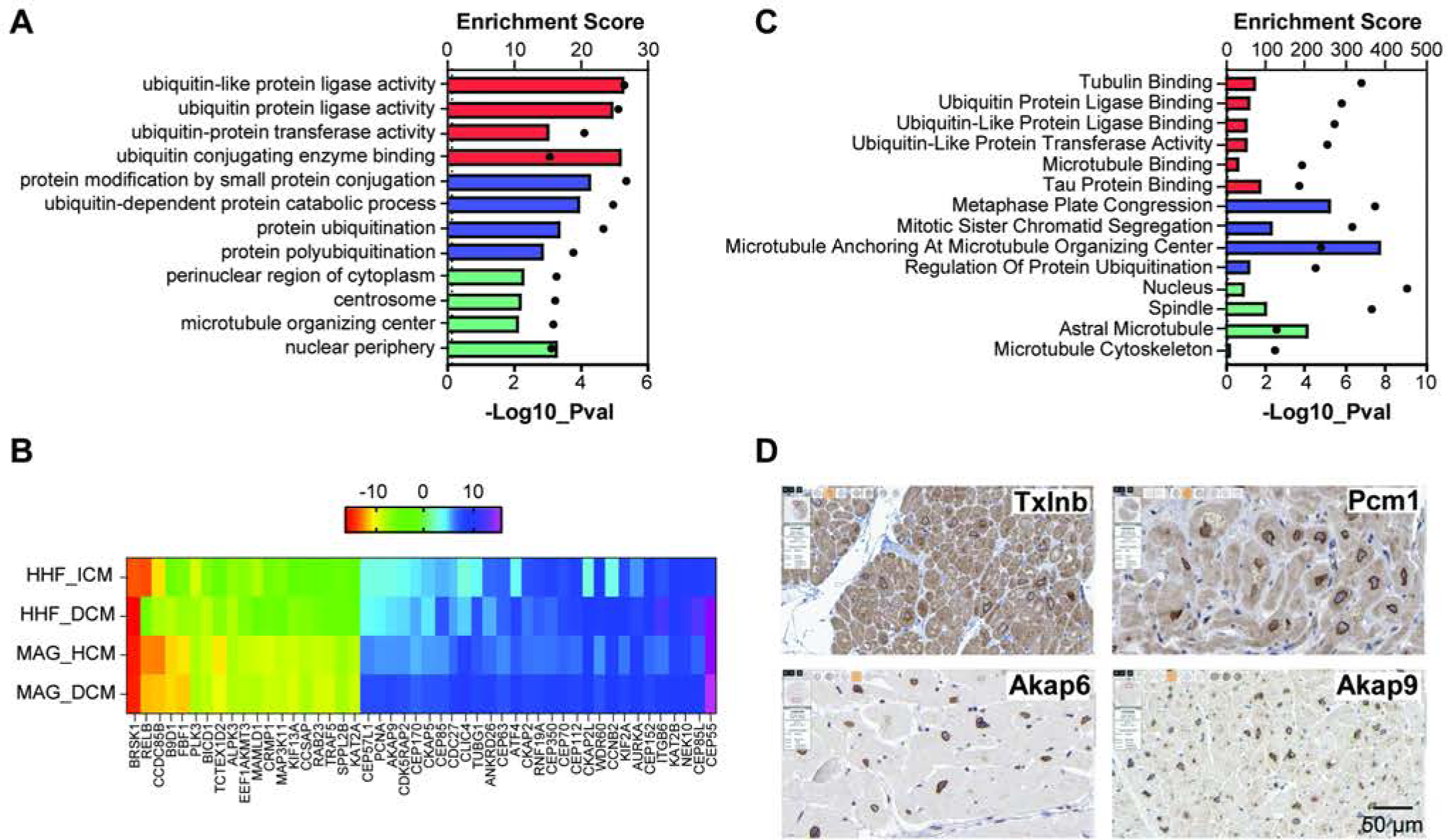
Centrosome-associated genes are broadly dysregulated in heart failure. (**A**) Enrichr gene ontology analysis for dysregulated genes in human heart failure enriches perinuclear, centrosome, MTOC, and ubiquitination pathways. The pathways are plotted as the enrichment score (bars, top axis) and negative log10_pval (dots, bottom axis) from the Fisher exact test and were selected from the top ten of each GO category, shown as bar color (red = Molecular Function, blue = Biological Process, and green = Cellular Component). Complete data in Supplement Table 2. (**B**) Heatmap of selected centrosome-associated gene expression significantly dysregulated (Adj. pval < 0.05) in at least three of four human heart failure RNAseq datasets, selected from a total of 43 upregulated and 128 downregulated genes. Public datasets are HHF-ICM (PMID30419824, GSE116250), HHF_DCM (PMID30419824, GSE116250), MAG_DCM (PMID36776621, GSE141910), and MAG_HCM (PMID36776621, GSE141910). Centrosome-associated genes are selected from ontologies: microtubule organizing center (GO:0005815), centrosome (GO:0005813), and centriole (GO:0005814). Color scale represents a normalized effect size calculated from a weighted Log2 Fold Change. Complete data in Supplement Table 3. (**C**) Enrichr gene ontology analysis for taxilin family protein interactors enrich for centrosome, microtubule, cytoskeleton, and protein ubiquitination pathways. Represented pathways are plotted as described above. Complete data in Supplement Table 4. (**D**) Immunohistochemistry in human heart tissue shows prominent perinuclear localization of Txlnb in non-failing human heart tissue similar to centrosome proteins Pcm1, Akap6, and Akap9 (images from The Human Protein Atlas Version 23.0).

Next, we interrogated the taxilin protein interactome using curated high throughput mass spectrometry data from the BioGRID database^22,23^ and found a broad taxilin interaction network consisting of about 300 proteins with many shared interactors across the three isoforms **(Supplement Table/Figure 1e)**. Gene ontology and pathway analyses clustered this broad taxilin interaction network into cytoskeletal, microtubule, and UPS categories, with a most prominent enrichment for centrosome-related terms **(Figure 1c, Supplement Table 4)**. Given that taxilins are understudied in the heart, we assessed their expression and corroborated that Txlnb is highly enriched in cardiomyocytes **(Supplement Figure 1f)**. We next examined Txlnb localization in skeletal and cardiac muscle by querying the Human Protein Atlas (HPA) database Version 23.0^24,25^. Within cardiomyocytes, Txlnb protein localizes distinctly to narrow perinuclear compartments (**Figure 1d)**, consistent with the location of many centrosomal proteins, including the well-established centriolar-associated proteins Pcm1, Akap6, and Akap9 that redistribute to form perinuclear microtubule “cages” in cardiomyocytes. Notably, perinuclear enrichment of cardiomyocyte proteasomes has also been previously observed by Gomes et al,^26^ (see figure 5 in cited reference) and is further supported by HPA immunohistochemistry images for several proteasomal proteins including Psma4, Psma5, Psmb1, Psmd1, among many others **(Supplement Figure 1g)**. Given GO enrichments of UPS and centrosomes and the co-localization of Txlnb, MTOC, and proteasomes, we speculate that Txlnb may provide a functional link between these organelles.

### 2.2 Overexpression of Txlnb improves proteasomal insufficiency during NRCM hypertrophy

Prior studies indicate that taxilin proteins are centrosomal adapters and that Txlng influences the accumulation of ubiquitinated proteins^27^, suggesting that taxilin proteins may act at the interface of the centrosome-proteasome axis. To begin examining this, we focused on establishing a role for Txlnb in cardiomyocyte proteostasis, while testing whether Txlnb overexpression is sufficient to modulate cardiomyocyte UPS function. Neonatal rat cardiomyocytes (NRCMs) were isolated and transduced with adenovirus encoding either Txlnb-GFP (C-terminal GFP Fusion), or GFP only, and stimulated *in vitro* with phenylephrine (PE) to induce hypertrophy accompanied by increased protein synthesis and accumulation of ubiquitin-conjugated proteins targeted for degradation. Compared to GFP controls, Txlnb overexpression significantly reduces the accumulation of ubiquitinated proteins **(Figure 2a-b)** and significantly increases β5 proteasome activity **(Figure 2c)** after PE treatment. Next, we performed puromycin labeling of nascent polypeptides to assess if Txlnb influences protein synthesis and found that Txlnb overexpression increases puromycin-labeled proteins **(Figure 2d-e)**. This unexpected observation is also supported by simple luciferase-based reporter assays where overexpression of Txlnb (native or GFP-tagged) is sufficient to increase reporter expression/activity at baseline, which is further exacerbated with PE treatment **(Figure 2f)**. Together, these data suggest that Txlnb regulates proteostasis in cardiomyocytes, potentially through coordinated regulation of protein synthesis and degradation through the UPS.

**Figure 2.**
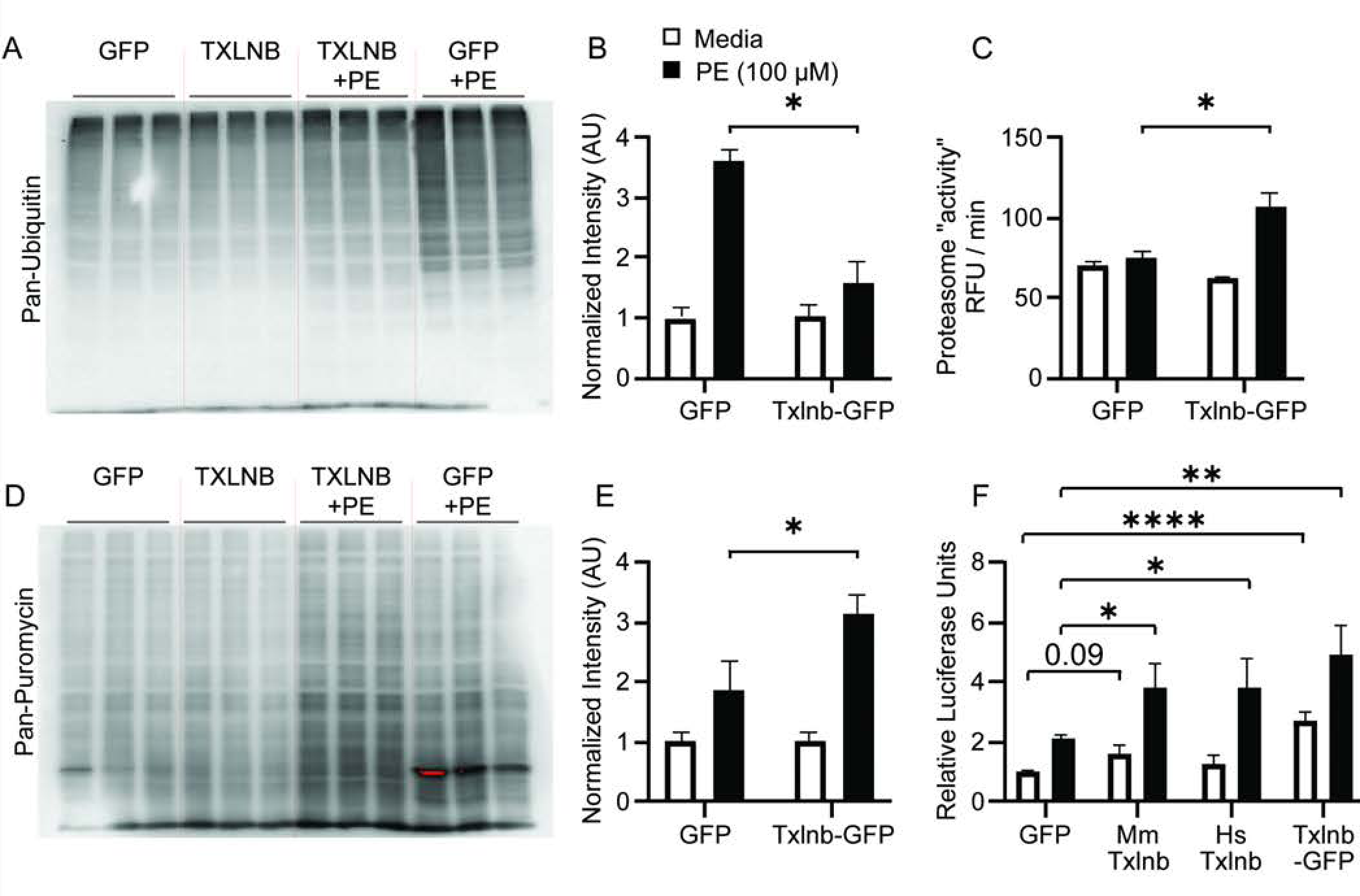
Overexpression of Txlnb improves proteasomal insufficiency during NRCM hypertrophy. (**A**) Western blot and (**B**) data quantification for ubiquitin conjugates from NRCM lysates infected with Ad5-GFP or Ad5-Txlnb-GFP, and treated with, without phenylephrine (PE, 100 μM) shows that overexpression of Txlnb reduces PE induced ubiquitin conjugates (n=6 per group, *=p<0.05). (**C**) Ad5-mediated overexpression of Txlnb in NRCMs increases 26s-β5 proteasome activity after PE treatment (n=5 per group, *=p<0.05). (**D**) Western blot and (**E**) data quantification for Sunset assay (puromycin labeling of new peptides) in NRCMs infected with Ad5-GFP or Ad5-Txlnb and treated with or without phenylephrine (PE, 100 μM) indicates that overexpression of Txlnb increases protein synthesis (n=6 per group, *=p<0.05). Statistics used for panels B, C, and E were one-way ANOVA, followed by Dunnet’s post-hoc test. (**F**) Protein abundance readout from luciferase reporter indicates that Txlnb clones from both mouse and human increase luciferase expression (n>6 per group, * = p<0.05, One-way ANOVA, followed by Sidak’s multiple comparisons testing GFP-PE vs each Txlnb-PE group). For panels B, C, E, and F, black bars indicate cells treated with 100 µM PE, whereas open bars are serum free media only.

### 2.3 Txlnb knockout mice show reduced cardiac mass with age but maintain cardiac function

The UPS system is required for cardiac homeostasis, and UPS dysfunction is implicated in heart growth, as well as the onset and progression of heart failure.^28^ To determine if Txlnb regulates cardiac structure and function, we developed germline Txlnb-knockout (KO) mice on the C57BL/6J background using Crispr-Cas9 technology with guide pairs flanking exon 3 to produce a deletion that would lead to mis-splicing (exon 2-4) and nonsense-mediated decay of frame-shifted mRNA. Indeed, Txlnb-KO mouse hearts show a specific and complete loss of Txlnb protein, relative to wildtype (WT) littermate control hearts **(Figure 3a),** as well as a 97% reduction in Txlnb mRNA **(Figure 3b).** Although the calculated molecular weight of Txlnb is 77 kDa, this protein routinely migrates at a molecular weight of 120 kDa^10,13^, consistent with positive control lysates that we generated with overexpression of mouse and human cDNA sequences. In addition, Txlnb-KO mouse heart tissue sections show a loss of Txlnb immunostaining, which in WT mice, appeared to predominantly localize to the perinuclear space in cardiomyocytes as expected **(Figure 3c).**

**Figure 3.**
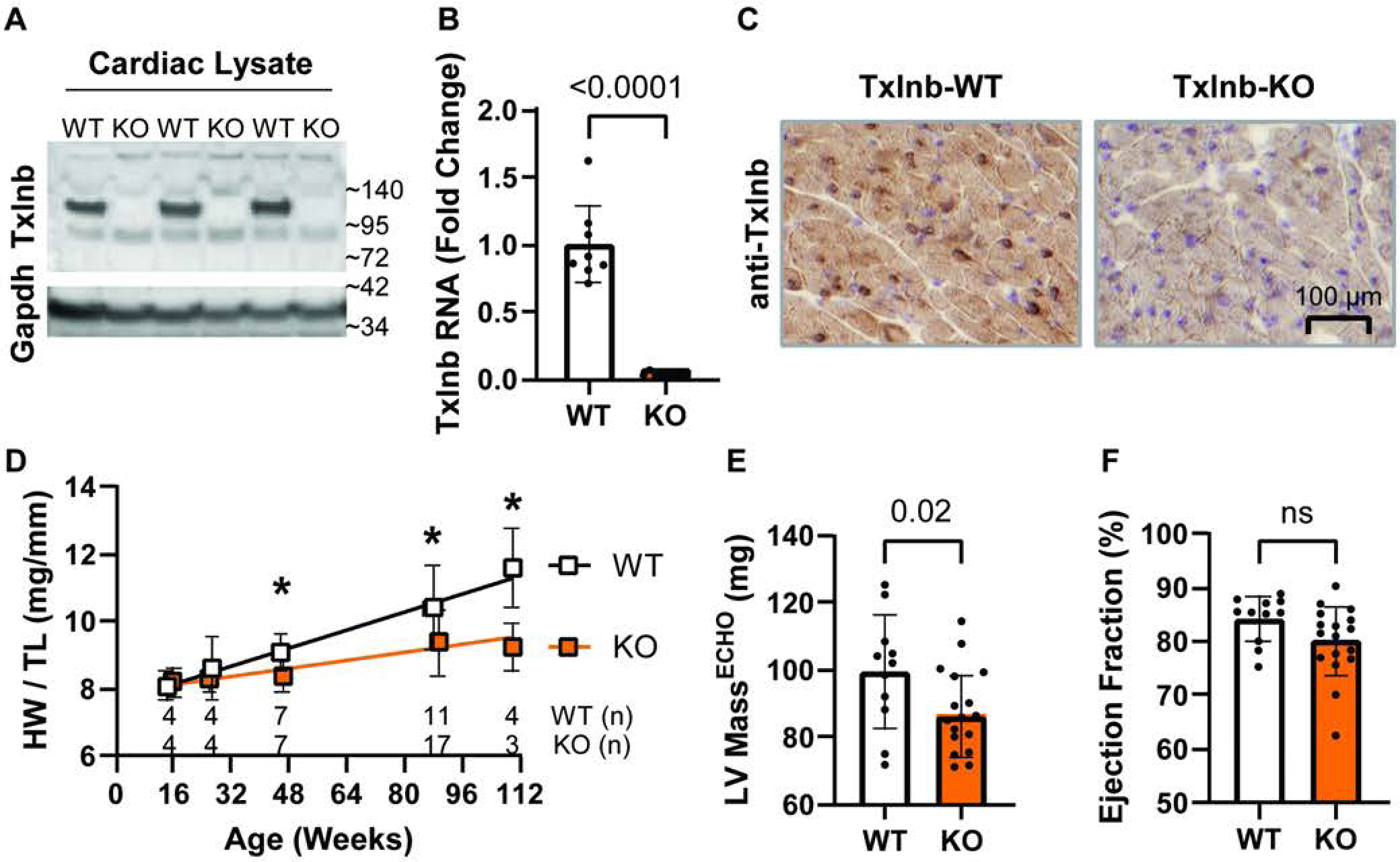
Txlnb knockout mice show reduced cardiac mass with age but maintain cardiac function. (**A**) Confirmation of Txlnb protein knockout in heart tissue using western blot analysis with antibody HPA030403. (**B**) QPCR results showing loss of Txlnb expression in KO mice. N=10 per genotype. P<0.0001, normalized to GAPDH via 2 ΔΔCt method. (**C**) Immunohistochemistry results showing prominent perinuclear localization of Txlnb in WT hearts and loss of Txlnb expression in KO hearts. (**D**) Intermittent gravimetric analysis of indexed heart mass (heart weight normalized to tibia length) showing that Txlnb deletion decreases age-related cardiac hypertrophy (n indicated on plot, *=p<0.05, individual T-tests, trendline from linear regression analysis). Echocardiography analysis at 18 months of age shows reduced heart mass in Txlnb knockout mice (**E**) and normal cardiac ejection fraction (**F**).

Txlnb-KO mice are viable, with normal postnatal development and adult maturation, and present no overt abnormalities. To determine the effects of Txlnb knockout on cardiac structure and function, we collected echocardiographic measures in mice up to ∼2 years of age and made closer examinations of mouse size, heart size, and heart weight (HW) normalized to tibia length (HW:TL ratio). We found that WT mice exhibit characteristic age-related increases in HW:TL, whereas Txlnb-KO mice do not increase cardiac size as they age **(Figure 3d)** (Difference in Slopes, WT=0.0335, KO=0.0154, pval=8.3e-3). This is further supported by echocardiography data collected at ∼18 months of age, indicating that Txlnb-KO mice have decreased cardiac mass compared to WT **(Figure 3e)**. Despite the smaller size, Txlnb-KO hearts maintain normal cardiac function, i.e., ejection fraction is not different **(Figure 3f).** Together, these data support our successful generation of global Txlnb-KO mice, which are mostly normal but display altered cardiac growth with aging.

### 2.4 Txlnb knockout mouse hearts exhibit signs of proteasomal insufficiency

Data from our *in vitro* studies show that overexpression of Txlnb is sufficient to increase proteolysis of ubiquitinated proteins through the UPS in cultured NRCMs. To determine if Txlnb is required for effective UPS function *in vivo*, we extracted and analyzed protein lysates from WT and Txlnb-KO mouse hearts harvested at 16 weeks of age and found that Txlnb-KO samples show a significant accumulation of ubiquitinated proteins (∼2-fold, P<0.0001) **(Figure 4a-d)**. Next, we assayed the activities of the three catalytic subunits (B1, B2, B5) of the proteasome and showed that Txlnb-KO hearts have significantly decreased B5 proteasome activity (∼25%, p<0.01) **(Figure 4e)**, while B1 proteasome activity trended upward (p=0.15), and B2 activity remained unchanged from a low basal rate. Together, these observations (increased ubiquitin-conjugate accumulation and decreased proteasome activity) are consistent with cardiac proteasomal insufficiency, which in the setting of cardiac stress, has been shown to contribute to significant cardiac dysfunction and premature death.^29^

**Figure 4.**
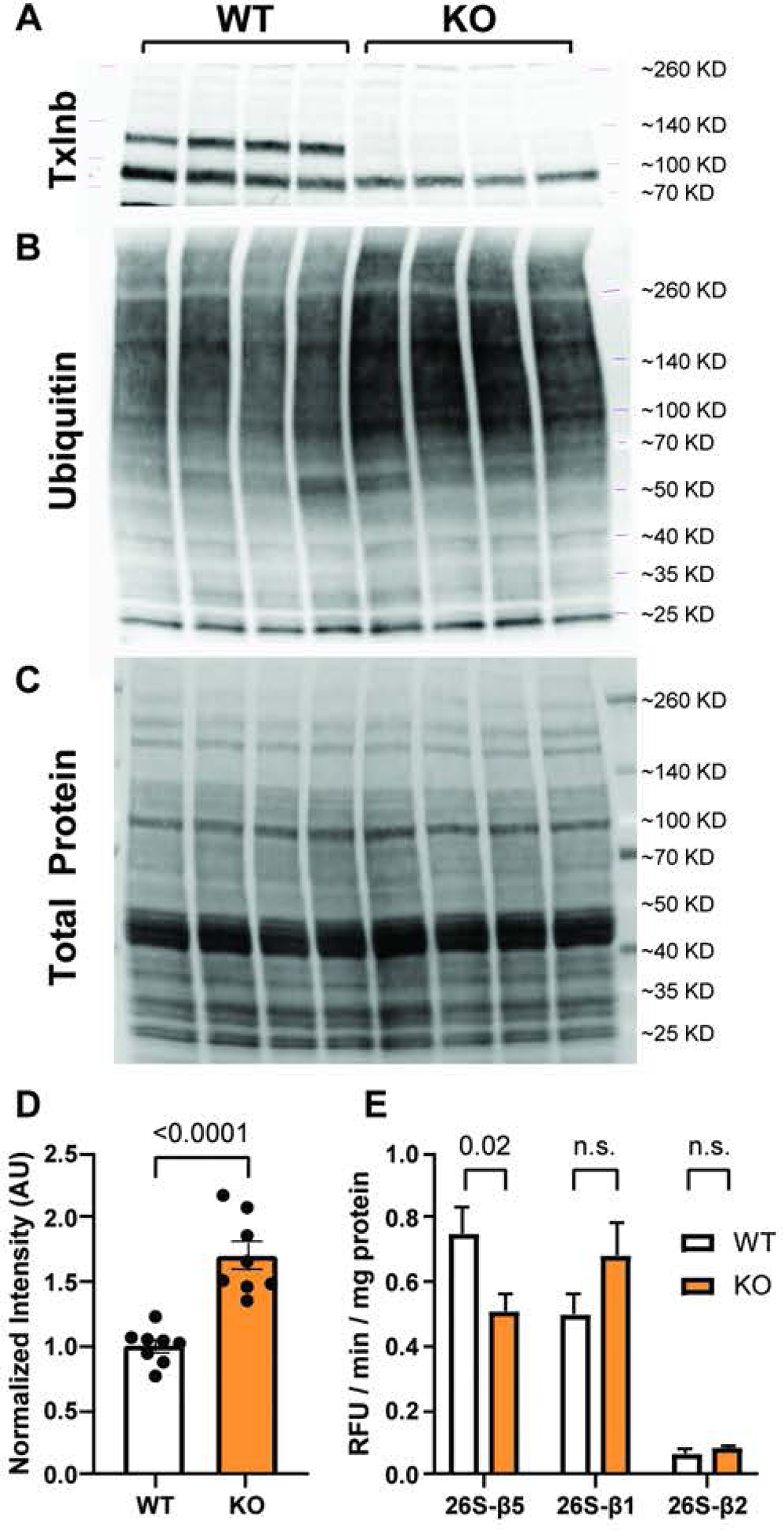
Txlnb knockout mouse hearts exhibit signs of proteasomal insufficiency. (**A-C**) Western blot for (A) Txlnb, (**B**) ubiquitin conjugates, and (**C**) total protein in mouse heart lysates. (**D**) quantification of ubiquitin conjugates blot normalized to total protein shown in B-C (n=8 per group, p<0.0001, unpaired two-tailed t-test). (**E**) 26S-proteasome activity assays for cardiac lysates specific for β-1, β-2, and β-5 activity in WT and Txlnb-KO heart lysates (N=8 per group, p-value determined by unpaired two-tailed t-test for each proteasome activity).

### 2.5 Loss of Txlnb worsens proteotoxic heart failure in Cryab-R120G transgenic mice

We next investigated if Txlnb loss influences heart failure caused by proteotoxic stress. We crossed Txlnb-KO mice to transgenic Myh6-Cryab-R120G mice, a well-established model of cardiac proteotoxicity causing significant heart failure and premature death^30^, and performed cardiovascular phenotyping between littermates. Echocardiography at 10 months of age shows the expected reduction in systolic function (ejection fraction, EF) caused by Cryab-R120G overexpression, and notably, this is further exacerbated in Txlnb-KO/Cryab+ transgenic mice (EF 45% vs 62%, p<1e-4) **(Figure 5a)**. This decreased EF was coincident with increased end systolic volumes **(Figure 5b)** in Txlnb-KO/Cryab+ transgenic mice, compared to WT/Cryab+ mice. Pericardial effusions were noted in 5 of 34 Txlnb-KO/Cryab+ mice, but only 1 of 35 Txlnb-WT/Cryab+ littermates **(Supplement Table 5)**. Heart tissues were collected and separated into soluble versus insoluble protein lysates for western blotting of ubiquitin-conjugated proteins. As expected, Cryab-R120G positive samples show an overwhelming accumulation of ubiquitinated proteins (primarily in detergent-insoluble fractions), which is slightly increased (p-value=0.0507) on the Txlnb-KO background **(Figure 5c-f)**. Together, these data suggest that loss of Txlnb worsens Cryab-R120G proteotoxic stress-induced heart failure in mice, perhaps through subtle influence on ubiquitinated protein accumulations given that cardiac detriments in these mice take months to manifest.

**Figure 5.**
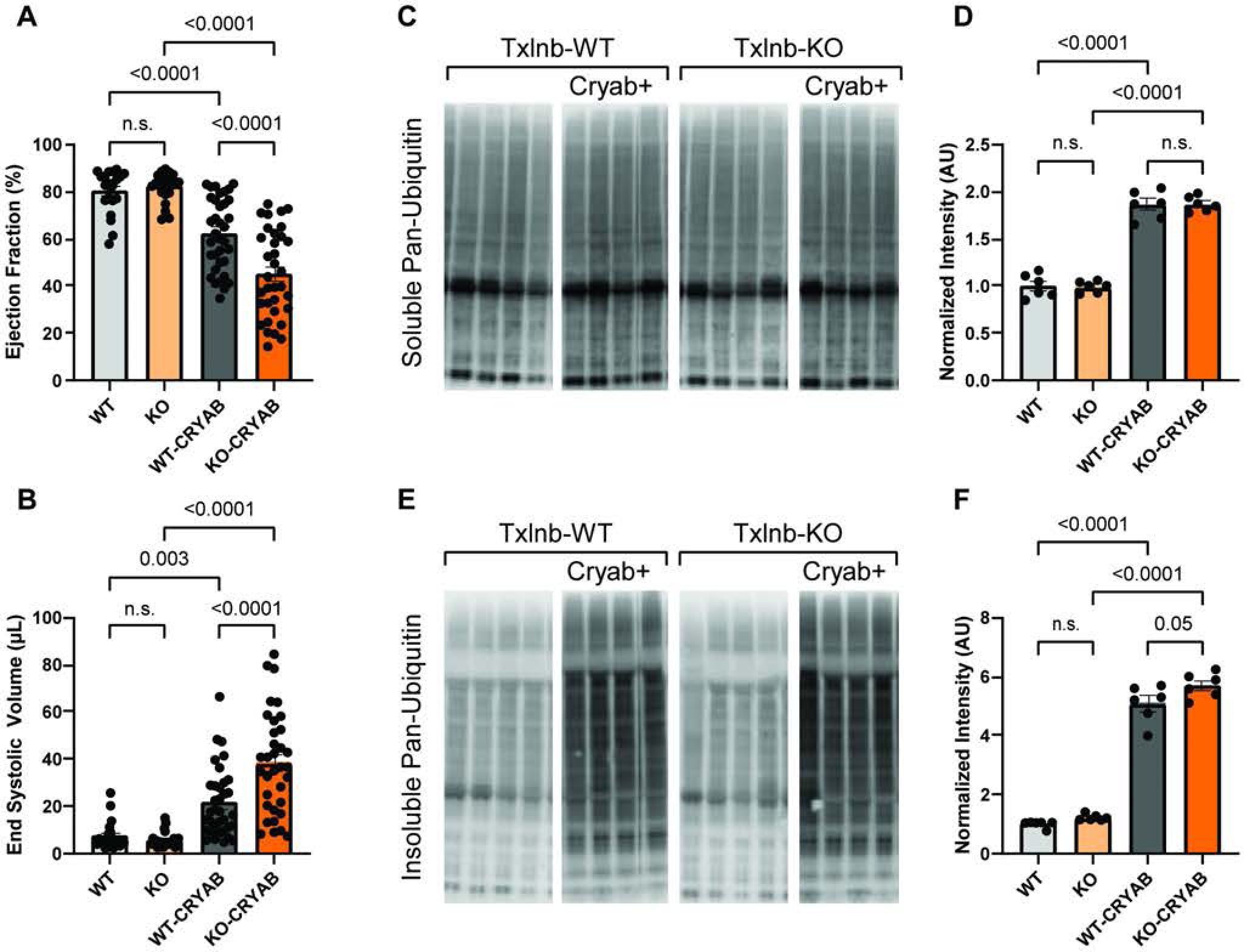
Loss of Txlnb worsens proteotoxic heart failure in Cryab-R120G transgenic mice. (**A-B**) Echocardiography at 10 months of age shows Txlnb-KO/Cryab-R120G+ mice have reduced ejection fraction (**A**) and increased end-systolic volume (**B)** compared to littermates (n=21-35 mice per genotype, p-value from one-way ANOVA with Sidak’s posthoc test). (**C-F**) Cardiac lysates were separated into detergent soluble (**C**) vs insoluble (**E**) fractions and analyzed for ubiquitinated conjugates via western blot and normalized to total protein from stain-free gel images (not shown). Quantified data is presented in panels D and F (n>6 per genotype, p-value from one-way ANOVA with Sidak’s posthoc test).

### 2.6 Loss of Txlnb exacerbates proteostatic perturbations during cardiac hypertrophy in mice

Our data suggest that Txlnb regulates proteostasis in mouse hearts and during cardiomyocyte hypertrophy in PE-treated NRCMs. We next sought to determine if loss of Txlnb exacerbates structural and functional remodeling in a mouse model of cardiac hypertrophy. WT and Txlnb-KO mice at 12 weeks of age were subjected to surgically-induced chronic pressure overload through a minimally invasive transverse aortic constriction (TAC). At 6 weeks post-surgery, echocardiography data show that minimally invasive TAC does not cause an obvious decrease in ejection fraction compared to sham-operated controls, which is typical for C57BL mice on the 6J background^31,32^ **(Figure 6a)**. However, female Txlnb-KO mice subjected to TAC did exhibit a significant reduction in EF compared to WT, which is interesting given that female mice (versus males) generally show more favorable responses to TAC. Mice were subsequently euthanized, and hearts were collected for morphological and molecular measures. Gravimetric analyses show that both male and female WT and Txlnb-KO mice have significant cardiac hypertrophy following TAC, however there is no difference in the response between genotypes **(Figure 6b).** Most notably, western blot analysis **(Figure 6c)** shows that TAC causes a significant accumulation of ubiquitin-conjugated proteins in WT mice as expected and that this is exacerbated in Txlnb-KO hearts **(Figure 6d)**. These data corroborate that loss of Txlnb disrupts proteostasis in mice by revealing a profound influence on ubiquitinated protein abundances in the setting of cardiac hypertrophy.

**Figure 6.**
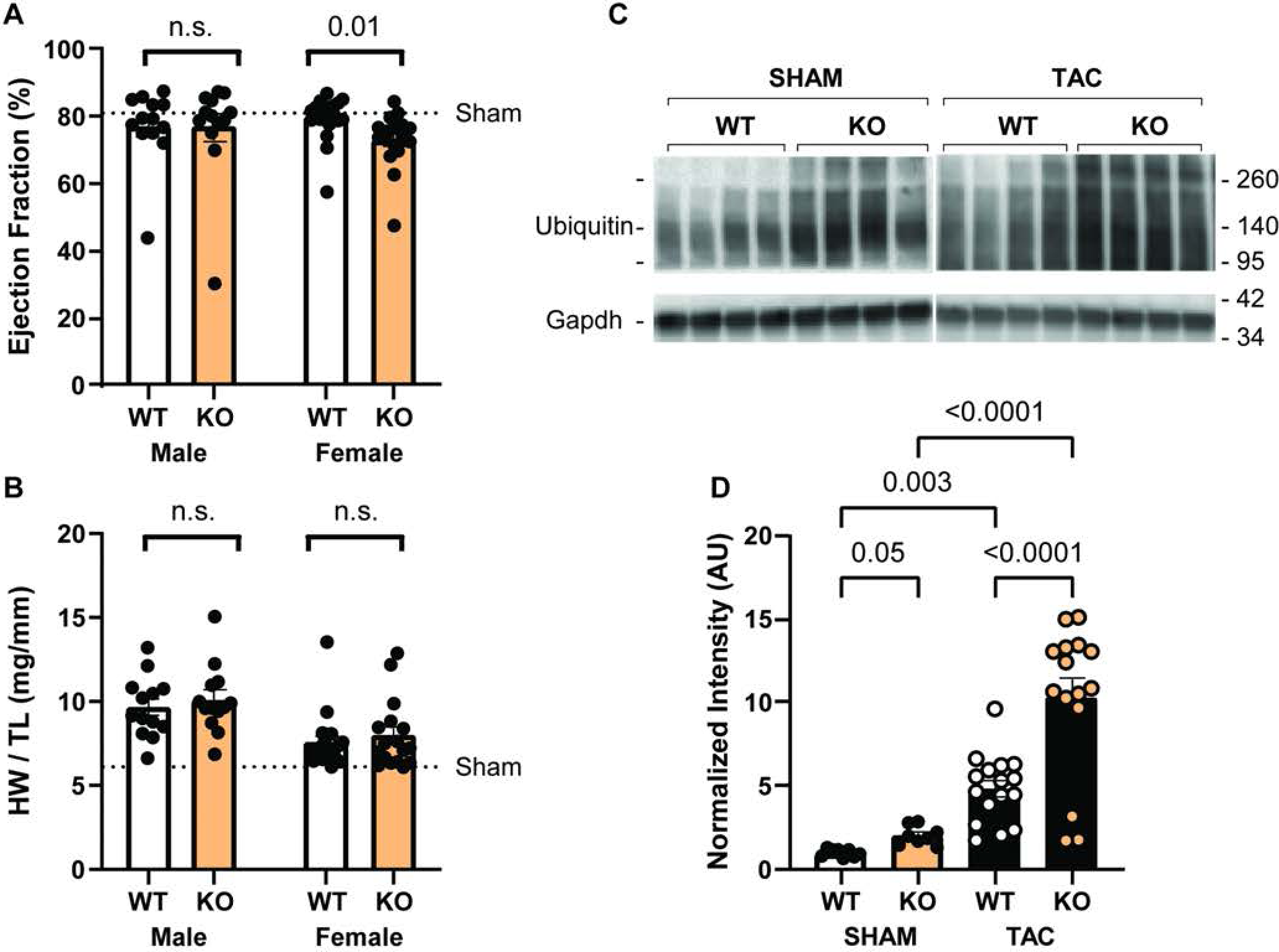
Loss of Txlnb exacerbates proteasome dysfunction during cardiac hypertrophy in mice. (**A**) LV pressure overload via TAC for 6 weeks caused a significant reduction in ejection fraction in female Txlnb-KO mice compared to WT littermates (n=17-20). Male Txlnb-KO mice were not different from WT littermates (n=13 each). P-value calculated using unpaired two-tailed t-test. (**B**) Post-mortem gravimetric analysis (heart mass normalized to tibia length) indicates that both WT and KO, Male and Female mice had cardiac hypertrophy after TAC, without a significant difference by genotype (n=13-20 per group, p-value from t-test between genotype by sex). (**C-D**) Western blot analysis shows an accumulation of ubiquitin conjugates in Txlnb-KO hearts that is exacerbated after LV pressure overload via TAC normalized to Gapdh, (n=8-16 per group, p-value from one-way ANOVA with Sidak’s posthoc test).

### 2.7 Loss of Txlnb contributes to dysregulated centrosomal protein expression

Our data show that loss of Txlnb causes proteasomal insufficiency in mice at baseline and can aggravate proteotoxic and hypertrophic cardiomyopathy in mice. To begin delineating the underlying mechanisms, we next assessed whether altered proteasome activity in Txlnb-KO mice might be related to altered expression of proteasome subunits (i.e. inferred proteasome abundances). Western blotting for several proteasomal proteins, including Rpt1 and PA28a (activator proteins) and Psmb1, Psmb2, and Psmb5 (catalytic proteins) in total cardiac protein lysates show that Txlnb deletion does not perturb proteasomal subunit protein expression **(Figure 7a-b)**. Notably, along with these blots, we also blotted for Txlna, which is the second most abundant taxilin protein in the heart and were intrigued to find increased Txlna (∼6-fold, p=1e-4) in Txlnb-KO hearts, perhaps reflecting a compensatory mechanism to maintain integrity and function of specialized centrosomes in cardiomyocytes. Considering that Txlnb-KO did not disrupt proteasomal protein expression but did ectopically upregulate expression Txlna, we next evaluated the expression of other centrosome proteins. Using a candidate-based approach (based on BioGRID associations, RNA co-expression with Txlnb, and known interactions with taxilin paralogues). We measured the expression of centrosomal proteins Txlng, Cep63, Ofd1, and Tubg via western blot analysis of WT and Txlnb-KO heart tissue **(Figure 7c)**. Western blot data show that Cep63 is significantly upregulated in Txlnb-KO hearts (∼2-fold, p=0.01). Notably, Cep63 was previously identified as a Txlnb interacting protein and its RNA is co-expressed with Txlnb in human hearts (r^2^∼80-90%, FDR<1e-5). Data also shows that knockout of Txlnb causes a small but significant increase in Txlng expression (p=0.036), a trending increase in Ofd1 (p=0.09), about a 2-fold trending increase in gamma tubulin expression (Tubg, p=0.07) **(Figure 7d)**. Together, these data provide evidence supporting that Txlnb may act primarily at the level of the centrosome, where it influences UPS function by means that remain unresolved.

**Figure 7.**
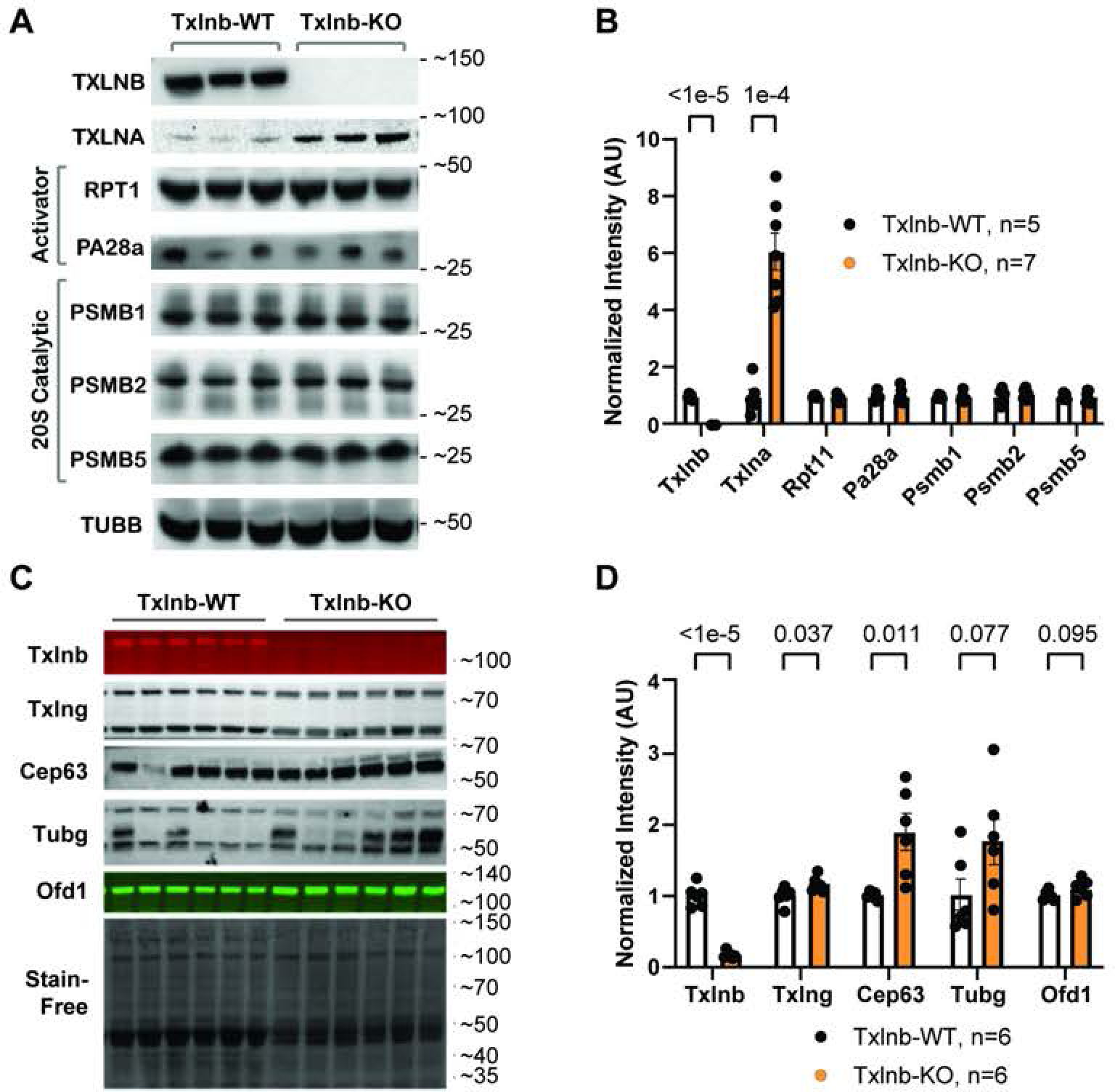
Txlnb-KO contributes to dysregulated centrosomal protein expression. (**A-B**) Western blot to assess protein expression of proteasome subunits in WT and Txlnb-KO cardiac lysates; note compensatory upregulation of Txlna, (n=5-7 per genotype, p-value from unpaired two-tailed t-test per protein target, normalized to Tubb). (**C-D**) Western blot to assess protein expression of centrosome proteins in WT and Txlnb-KO cardiac lysates; note upregulation of Cep63 and Tubg (n=6 per genotype, p-value from unpaired two-tailed t-test per protein target, normalized to Stain-free).

### 2.8 Txlnb model

Our data suggest that the centrosome-associated protein Txlnb regulates proteostasis in cardiomyocytes through an unknown molecular mechanism. Structure predictions based on Txlnb protein domains show intrinsically disordered N- and C-terminal domains (IDPR) that flank a large coiled-coil domain (CCD)) that may facilitate protein-protein interactions **(Figure 8a)**. Structural modeling and other experiments suggest that Txlnb, along with other taxilin proteins, lacks enzymatic activity related to ubiquitin ligation and proteolysis. Thus, we speculate (based on BioGRID interactome and localization data) that Txlnb anchors to the redistributed perinuclear centrosomal MTOC and interacts with diverse UPS-targeted proteins via its IDPRs and CCDs to improve their delivery to proteasomes. Using software (iTASSER^33^ and PHYRE2^34^) we developed a consensus model that shows profound similarity across taxilin proteins **(Figure 8b)**. The models show a “Y” like structure with the N- and C-terminal IDPRs forming two “arms”, the conserved CCD folding into a stable “stalk”, and the distal “foot” that may resemble a microtubule-binding domain, as PHYRE2 indicated structural homology between taxilin proteins and dynein, a classic microtubule-binding motor protein. In addition, we found that perinuclear localization of Txlnb in NRCMs is microtubule dependent (i.e. disrupted by classic microtubule depolymerizing agents nocodozole and colchicine) **(Figure 8c)**. In conclusion, we speculate that Txlnb functions as a specialized microtubule-dependent centrosomal-adapter protein that functionally bridges perinuclear centrosomes with transient and dynamic interactions with the proteasome or ubiquitin substrates to promote their delivery to proteasomes **(Figure 8d)**.

**Figure 8.**
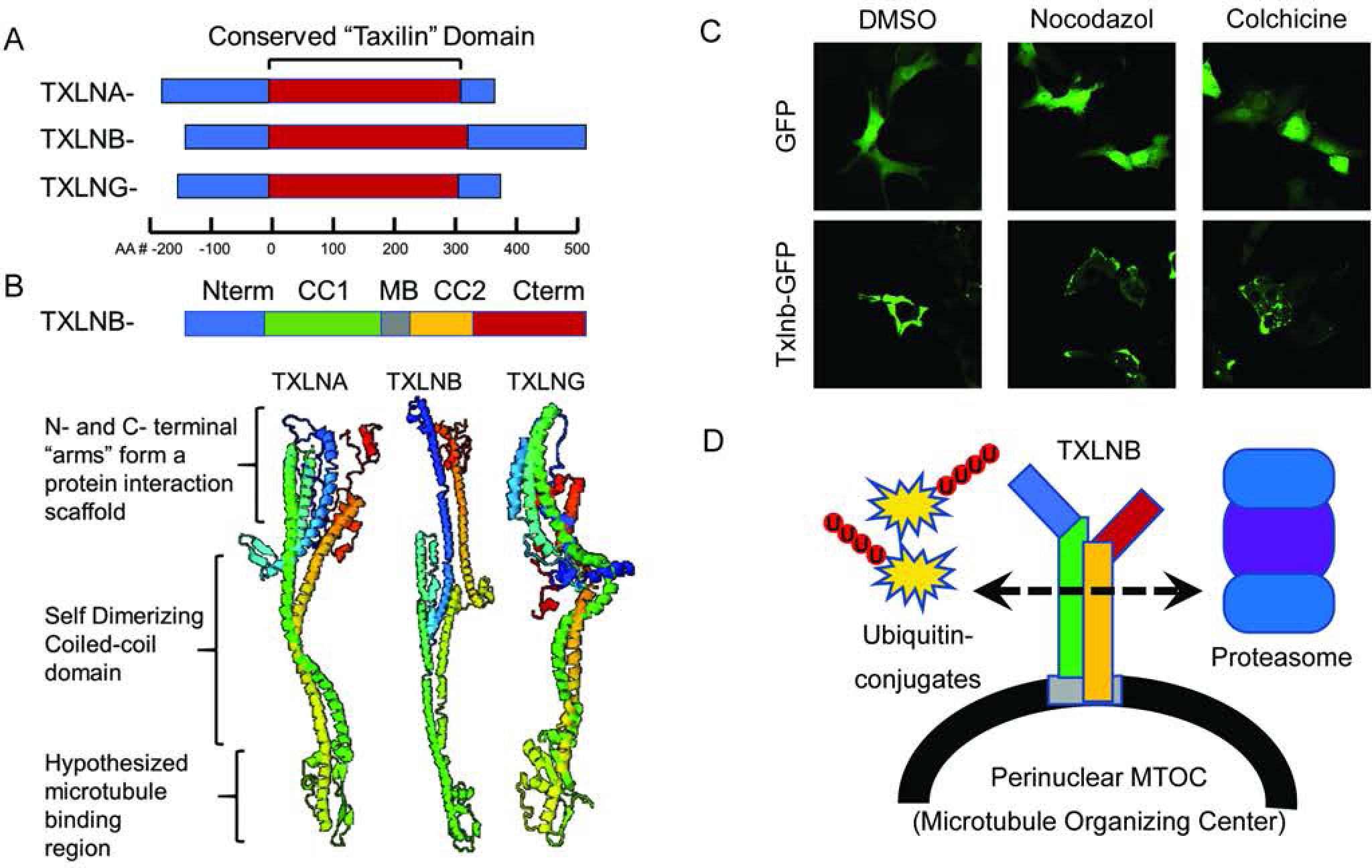
Txlnb may function in cardiomyocytes as a specialized microtubule-dependent centrosomal-adapter protein to regulate the UPS. (**A**) Primary sequence alignment among taxilin paralogues, indicates the large conserved coiled-coil domain (taxilin domain, colored red), flanked by variable length intrinsically disordered regions on the N- and C-termini (colored blue), note the absence of other predicted enzymatic or catalytic domains. (**B**) Computational model of Txlnb secondary structure. Note the predicted microtubule-binding domain (colored grey) centrally located between two coiled-coil domains (colored green and yellow). The ribbon diagram is colored N-to-C with annotations depicting a potential microtubule-binding domain predicted by PHYRE2. (**C**) Intracellular location of GFP or Txlnb-GFP transduced in NRCMs shows perinuclear localization of Txlnb is disrupted after 3-hour treatment with microtubule polymerization inhibitors Nocodazole (10 µM) or Colchicine (1 µg/mL). (**D**) A proposed model where Txlnb is anchored to the perinuclear MTOC and functions in a specialized centrosome-associated UPS.

### 3.0 Discussion

This study began with our interest in understanding the biological relevance of centrosomes in the heart. This was inspired by observations that the expression of centrosome-associated genes is profoundly dysregulated in heart failure, despite the absence of prototypical centrosome structures in cardiomyocytes. While exploring the relationship between centrosomes and heart failure, we identified Txlnb which belongs to the taxilin family, an understudied group of centrosome-adapter proteins. To further investigate the role of Txlnb in cardiac biology, we first focused on Txlnb functions in cardiomyocytes, where it is highest expressed, and performed a series of complementary gain- and loss-of-function studies in both cells and mice to decipher its causative effects. The results overwhelmingly support a role for Txlnb in influencing centrosomal composition and regulating cardiomyocyte proteostasis through the ubiquitin-proteasome system (UPS).

The UPS is responsible for degrading most intracellular proteins^35^ and exhibits consistent dysregulation in failing human hearts^36–38^. Furthermore, UPS function is a key determinant of HF outcomes in mice^28,39^. For example, myocardial damage from ischemia/reperfusion (I/R) injury is associated with UPS dysfunction, which causes the accumulation of diverse species of ubiquitinated and misfolded proteins and contributes to the molecular mechanisms of HF^39^. In addition, inhibiting proteasome function exacerbates myocardial IR injury^40^ increasing proteasome function protects against I/R injury^41^. It is interesting to note that in our studies, a two-fold accumulation of ubiquitinated proteins in Txlnb-KO mouse hearts was not sufficient to cause systolic dysfunction, even after ∼ 20 months of age. Even much more robust accumulations in Txlnb-KO TAC mice did not grossly perturb cardiac function; however, this may relate to the relatively short 6-week duration of our TAC studies. Longer-term studies should be carried out to assess if Txlnb-KO mice subjected to TAC develop cardiac dysfunction, as observed in Cryab-R120G mice after several months. Another consideration for our observations is that recent work has highlighted localized proteostasis in cardiomyocytes with regional translation and protein degradation occurring at discrete spatial locations^42,43^. While there is a large pool of sarcomere-associated proteasomes, an equally large pool of perinuclear proteasomes exists^26^. Future work will need to examine the contribution of perinuclear versus sarcomeric UPS functions in cardiomyocytes and if and how accumulations of differential pools of ubiquitinated protein substrates impact cardiac function and stress responses. Given the predominant localization and proximity of Txlnb to perinuclear proteasomes and evidence of centrosome to proteasome crosstalk in non-myocytes, we speculate that Txlnb mediates a specialized centrosome to proteasome axis in cardiomyocytes that regulates perinuclear proteostasis.

In non-myocytes, bidirectional signaling between centrosomes and UPS supports a cooperative proteostatic signaling nexus^2^; however, canonical centrosomes do not exist in cardiomyocytes. Instead, a subset of centrosomal and adaptor proteins (such as Pcnt and Pcm1) are redistributed perinuclear during centrosomal reduction promoting terminal differentiation of cardiomyocytes^4,8^. Interestingly, centrosome reduction is a central part of spermatogenesis and Txlna and Txlng are implicated in testicular biology and fertility^44–46^. A recent publication implicates impaired centrosome reduction to cause infantile dilated cardiomyopathy, through loss-of-function mutations in rotatin (Rttn)^9^. Beyond this, much of the consequences of centrosomal reduction in cardiomyocytes have been focused on the reorganization of the MTOC, or terminal exit of cell-cycle proliferation^6–8^. Our results suggest that centrosome-related genes/processes are disrupted during heart failure, likely indicating functions beyond centrosome reduction or microtubule disorganization. This has the potential to expand the cardiac field’s focus to include other non-canonical areas of centrosome biology including the association with proteasomes. How the centrosomes and proteosomes coordinate within cardiomyocytes is an unexplored area of biology, and the taxilin family may be one functional crosslink that regulates perinuclear proteostasis.

Txlnb, Txlna, and Txlng are paralogues with no known catalytic activity^10^. Their sequences share a conserved internal coiled-coil domain (CCD) yet have unique N- and C-terminal intrinsically disordered protein regions (IDPRs). CCDs can form tight helical structures with high-affinity binding to other protein helices (e.g. other cellular CCD-containing proteins^47,48^). Indeed, the centrosome and pericentriolar material are composed of many CCD-containing proteins forming a disorganized matrix described as a sticky cloud-like amorphous structure^49,50^. IDPRs lack defined structures and instead are flexible to enable dynamic conformations and molecular contacts given the environment^51,52^. IDPRs have been implicated in protein chaperoning, scaffolding, and UPS function^53^. We speculate that taxilin family proteins have a similar ability to engage diverse sets of proteins through their CCD and IDPRs. While Txlnb is selectively expressed in striated muscle and has an unclear biological function, Txlna and Txlng are ubiquitously expressed across tissues, associate with centrosome and pericentriolar proteins ^11,12^, bind to microtubules^54^, regulate protein trafficking^55^ and influence the accumulation of UPS-targeted substrates^27^. Our results show that Txlnb increases UPS efficiency, though it remains unclear exactly how this occurs. While Txlnb was not identified in proteomic screens of cardiac 20S and 26S proteasome-associated proteins,^26,56^ and is not strongly associated with ubiquitin ligase complexes, perhaps due to being understudied. However, looking beyond centrosomal associations, the BioGRID interacting data among all taxilin proteins identifies several interesting, cardiac-relevant proteins including those related to ubiquitin ligation (Uba1, Ube2i, Cul3, Usp3, Ufd1, and Arih1), proteasomes (Psma4, and Psme2), and other UPS regulatory proteins (see Supp. Table 4), perhaps indicating multifaceted roles for taxilins in perinuclear proteostasis, and perhaps beyond, which could be a reflection of our proteasome activity assay data.

While our data suggests that Txlnb may influence a specialized centrosome to proteasome axis in cardiomyocytes; several outstanding questions remain, including the specific role of Txlnb and the unexpected compensation by the taxilin family. We speculate that Txlnb is anchored to the specialized centrosome through a direct interaction between the perinuclear MTOC and an internal microtubule-binding domain inside the “taxilin” domain. This model assumes a self-dimerizing coiled-coil domain to form a stable stalk-like structure and dynamic interactions with cellular substrates and cargo mediated through intrinsically disordered regions at the N- and C-termini. To rigorously examine this model will require extensive molecular phenotyping in a relevant cardiomyocyte model. Also, since Txlna and Txlng were unexpectedly upregulated in Txlnb-KO hearts, taxilin paralogues might have compensatory and redundant roles in cardiomyocytes and might explain why cardiac phenotypes in Txlnb-KO mice are somewhat subtle. In addition, the unexpected sexual dimorphism that we observed in Txlnb-KO mice after TAC suggests males may be better protected than females, through unknown compensatory mechanisms. Without a better understanding of the regulation or molecular functions of taxilin paralogues in cardiomyocytes, it is difficult to predict how compensation could occur; however, the significant overlap of taxilin structures, interactors, and biological functions supports redundant molecular roles. Thus, future studies will need to test the effects of knocking out the entire taxilin family in cardiomyocytes using conditional approaches or begin to explore the role of each taxilin through AAV overexpression, with future potential as a gene therapy approach to treat proteasomal insufficiency for heart failure.

In summary, our studies demonstrate that the genetic knockout of the centrosomal protein Txlnb disrupts cardiomyocyte proteostasis, while Txlnb overexpression improves it. It is evident that Txlnb and taxilin paralogues serve vital roles in cardiomyocytes, extending beyond their reported influence on intracellular vesicular trafficking^10,57,58^. Significantly, our work validates previous studies that underscore the crucial role of centrosomes in mediating cardiomyocyte biology and expands the discussion to encompass their non-canonical roles, including the regulation of perinuclear ubiquitin-proteasome system. Our findings carry implications for cardiac aging and heart failure, particularly in the context of cardiac proteostasis. We emphasize the necessity for additional research to elucidate both the global and spatial mechanisms governing cardiac proteostasis, which includes sarcomere-associated, centrosome-associated, and potentially other subcellular regions.

---

**Supplement Figure 1.** Centrosome dysfunction underlies heart failure and may involve Txlnb. (**A-C**) Centrosomal associations between taxilin paralogues and centrosome proteins from a public BioID interaction study (PMID31304627) showing (**A**) Txlna association, (**B**) Txlng associations, and (**C**) Txlnb associations. Fold enrichment is represented with a bar graph on the left y-axis, and FDR is represented with dots on the right y-axis, where FDR<0.1 is shown in red. (**D**) RNA expression (TPM) of taxilin paralogues in HEK293 cells (from the human protein atlas). (**E**) BioGRID interactions for overlap of taxilin interacting proteins. (**F**) Txlnb RNA expression across human tissues shows muscle enrichment (from the GTEX consortium). (**G**) Immunohistochemistry in human heart tissue shows prominent nuclear/perinuclear localization of selected proteasome proteins Psma4, Psma5, Psmd1, and Psmb1 (images from The Human Protein Atlas Version 23.0).

## 5. Materials and Methods

### 5.1 Generation of Txlnb knockout mice

Germline Txlnb knockout mice were generated using CRISPR/Cas9 with assistance from the University of Iowa Genome Editing Core Facility. A pair of custom guide RNAs were designed to target intronic sequences flanking exon 3 of Txlnb. The upstream guide sequence is GGAGGGAACTTGCTGAGGCTCGT, and the downstream guide sequence is GAGAAGTGCACTGCGAACTGTGT. CRISPR components for pronuclear injection were prepared using previously described methods^59^. Briefly, spCas9 mRNA was *in vitro* transcribed after adding a T7 promoter to spCas9 coding region by PCR amplification of pX330. The two functional sgRNAs flanking Txlnb-Exon-3 were *in vitro* transcribed after generating templates with T7 promoters. SpCas9 mRNA and sgRNAs were purified using a MEGAclear kit (Life Technologies). The injection mix consisted of 100 ng/ml spCas9 mRNA and 50 ng/ml each sgRNA. Zygotes from 4 week-old B6SJLF1/J mice (The Jackson Laboratory Stock: 100012) were prepared for pronuclear injection^59^. Resulting offspring were genotyped to idenfity mice carrying the targeted Txlnb deletion, and the Txlnb-KO allele was then backcrossed to the C57BL/6J strain (The Jackson Laboratory Stock: 000664) for more than three generations prior to phenotyping. Size exclusion PCR was used for genotyping to detect WT from KO alleles. A second PCR reaction utilized a forward primer inside exon 3 to confidently identify Txlnb heterozygous from homozygous deletion mice. Primers used for genotyping are listed in **Supplement Table 6**. Four male transgenic Myh6-Cryab^R120G^ mice^30^ were acquired from Jeffrey Robbins (University of Cincinnati). Cryab mice were bred with Txlnb-/- females to establish F1 Txlnb heterozygous Cryab-positive mice. For subsequent breedings, the Myh6-Cryab^R120G^ transgene was carried with male breeders that are also heterozygous for Txlnb. Female breeders were heterozygous for Txlnb and Cryab Negative. Genotyping primers are listed in **Supplement Table 6**.

### 5.2 Transverse Aortic Constriction (TAC)

Minimally invasive TAC was performed as previously described with few changes.^60^ At the level of the suprasternal notch, a partial sternotomy and thyroid retraction were used to visualize the aortic arch on 8-week-old C57BL/6J male mice anesthetized with ketamine/xylazine (100/10 mg/kg, IP). The aorta was isolated and then constricted with a titanium ligating clip (Teleflex, #005200) gapped on 38-gauge acupuncture needles, and placed between the right innominate and left common carotid arteries. Sham mice underwent the same procedure but without constriction of the aorta.

### 5.3 Electrocardiography (EKG)

Mice were anesthetized with 2% isoflurane and placed on a surgical monitoring board to collect EKG and maintain body heat (Indus, Rodent Surgical Monitor+). Three-lead EKG was acquired using external titanium leads placed subcutaneously on limbs. Signals were routed through an analog output device (Indus, #351-0066-01) and converted into digital signals for analysis (AD Instruments, Powerlab 8/8, Labchart Pro v8.1.13, EKG module v2.4, and HRV module v2.0.3). Time selection windows of 10 seconds were used for EKG analyses with 4 beat averaging, and 60 seconds for HRV analyses, with standard mouse detection settings, and without excluding “outlier” beats. Data were exported for offline analyses using Microsoft Excel and GraphPad Prism v8.2.1.

### 5.4 Echocardiography

Echocardiography was performed on conscious mice, with mild sedation (Midazolam 5 mg/kg, SC) using a Vevo2100 imaging system (VisualSonics, Toronto, Canada). Mice were restrained in the left lateral recumbent position in the operator’s hand. Two-dimensional, M-mode, and Doppler imaging were performed in both long- and short-axis views to acquire standard measurements of left ventricular structure and cardiac function according to the endocardial and epicardial area protocol.

### 5.5 Western Blot

Mouse hearts were dissected, sectioned on a transverse plane, and frozen liquid nitrogen. The apical portion was homogenized in tissue lysis buffer (**Supplement Table 6.**) using a bead mill (Qiagen, TissueLyserII). Homogenates were sonicated, clarified by centrifugation (16,000×g, 10 mins, 4℃), and normalized by concentration using the BCA protein assay (Pierce, 23225). An equal mass of protein was separated via polyacrylamide gel electrophoresis using Criterion TGX Stain-Free midi gels (Biorad 5678084 or 5678085) or NuPAGE Bis-Tris gels (ThermoFisher, NP0336BOX), wet transferred to PVDF, (0.2 μm pore, Biorad), blocked in 5% NFDM in TBST, and incubated with primary antibodies diluted in 5% NDFM in TBST, overnight at 4℃. After washing, blots were incubated with species-specific secondary antibodies for 90 mins at room temperature. Antibodies are listed in **Supplement Table 6.** After washing, blots were imaged using Radiance Plus ECL substrate (Azure Biosystems #AC2103) or fluorescence on a Biorad Versadoc MP, or ThermoFisher iBright imaging system. Quantification of band intensity was done within the Biorad QuantityOne Software (Version 4.6.6) or iBright Analysis Software (version 5.0.1) with global background subtraction settings. Data were exported for offline analyses using Microsoft Excel and GraphPad Prism v8.2.1.

### 5.6 Tissue Immunohistochemistry

Mouse hearts were dissected, sectioned on a coronal plane, and frozen in OCT medium. Tissue blocks were sectioned at ∼5 micrometers, affixed to a glass slide, and postfixed in fresh 4% paraformaldehyde for 10 min at 4℃. Sections were permeabilized using 0.2% TritonX-100 in TBS and incubated with 3% hydrogen peroxide in water to quench endogenous peroxidase activity. Slides were blocked in 10% goat serum in TBS and incubated with primary antibodies diluted in blocking buffer (see table), overnight at room temperature in a humidified chamber. After washing, slides were developed using the protocol for the ABC-HRP Elite Rabbit-IgG kit (Vectastain, PK-6101) and Immpact DAB EqV (Vectastain, SK-4003). After washing, slides were counterstained in hematoxalin, washed, dehydrated, and mounted in Tek-Select Clear Permaslip Mounting Media (25% acrylic resin in toluene), coversliped, and imaged using an Olympus IX70 inverted fluorescence microscope with the 40x objective with a DP70 camera and DP Controller 2.1 software.

### 5.7 Sunset Assay

Assays were performed similarly to as described^61–63^. Mice were injected IP with 0.04 μmol/g body weight of puromycin (Corning Cellgro puromycin dihydrochloride #61-385-ra) dissolved in normal saline. After 30 mins, mice were euthanized, tissues were dissected, and snap frozen in liquid nitrogen. Powdered tissue was homogenized in tissue lysis buffer, quantified, and subjected to standard western blot protocols to detect puromycylated nascent polypeptides with an anti-puromycin antibody (DSHB #PMY-2A4). For in vitro sunset assays in isolated NRCMs, puromycin was added to cells at a final concentration of 1μM, and cells were incubated for 30 mins, before lysis in RIPA buffer and subsequent western blot.

### 5.8 Proteasome Assay

Assays were performed similarly to as described^61,64^. Tissues were homogenized in a proteasome activity assay buffer pH of 7.5 containing 50 mM Tris, 1 mM EDTA, 150 mM NaCl, 5 mM MgCl2, and 0.5 mM DTT added fresh. Tissue was powdered in liquid nitrogen, and homogenized in buffer using a bead beater (Qiagen TissueLyserII), at a ratio of ∼ 10:1 buffer to tissue. Lysates were centrifuged at 12,000 ×g for 30 mins at 4℃. The protein content of supernatants was quantified using the BCA method, samples were diluted to 2 mg/ml, and aliquoted for storage at −80℃. Specific proteasomal substrates and inhibitors that were used are listed in **Supplement Table 6**. Proteasomal activities were assayed with 10μg of protein and 50μM of AMC-labeled substrate in a total volume of 100μl. The 26S proteasome activities were measured in homogenization buffer with the addition of 100μM ATP and with or without the proteasome inhibitor Bortezomib at a final concentration of 2 mM. 26S proteasome activity assay was measured in triplicate, per condition fluorescent units from inhibitor-treated wells were background subtracted. The fluorescence of released AMC was quantified on a Spectramax M2 96 well plate reader with ex380/em460 filter settings in a kinetic assay at 15-minute intervals over 2 hours.

### 5.9 Cloning and Virus production

Several custom clones were used in this study. Txlnb was cloned from mouse heart cDNA using primers listed in **Supplement Table 6**. This was restriction enzyme digested and ligated into an AAV vector with a CMV-MCS-Ires-GFP (Uiowa AAV-939). Txlnb was amplified from this plasmid via PCR with primers F and R and subcloned into an Adenoviral production compatible plasmid (Uiowa Viral Vector Core) as a Txlnb-C-terminal GFP fusion protein (Txlnb-GFP). A plasmid encoding the human Txlnb sequence was purchased from the DNASU plasmid repository (clone HsCD00443576, pLX304 vector with a C-terminal V5 tag).

### 5.10 RNA Isolation

RNA was isolated using the “miRNeasy RNA isolation kit” (Qiagen, Ref# 217084). Sample lysates were prepared in 700 μl of Qiazol and stored at −80℃. Samples were thawed, incubated at room temperature for 5 mins, transferred to “Phasemaker tubes” (ThermoFisher Scientific, A33248), mixed with 140 μl of chloroform, and then centrifuged at 16,000 ×g for 15 mins. The upper aqueous layer was removed, combined with 550 μl of 100% ethanol, and loaded onto a purification column where a centrifuge or vacuum manifold was used to remove flow through. To wash columns, 350 μl of kit wash buffer RWT was washed through the column. To remove genomic DNA contamination, on column DNase digest was performed with the “RNase-Free DNase Set” (Qiagen, Ref# 79254). Lyophilized DNAse was resuspended in 500 μl of H2O, then each column received an 80 μl reaction (10 μl of DNAse with 70 μl of RDD buffer), and columns were incubated at room temperature for 15 mins. Next columns were sequentially washed with 350 μl of RWT buffer, 500 μl of RPE wash buffer, and 500 μl of RPE wash buffer using a centrifuge or vacuum manifold to remove flowthrough. Columns were transferred to an empty tube and recentrifuged at 8,000 × g for 1 min to remove all traces of wash buffer. Columns were transferred to an empty tube, 35 μl of nuclease-free water was added to the column membrane, and columns were incubated at room temperature for 1 min, then centrifuged at 8,000 rpm to elute RNA. RNA quality and concentration were determined using the “Qubit RNA High Sensitivity Assay Kit” (ThermoFisher Scientific, Ref# Q32852), and a nanodrop spectrophotometer.

### 5.11 Q-RT-PCR

For gene expression analysis, 1000 ng of total RNA was converted to cDNA using the “High-Capacity cDNA Reverse Transcription Kit” (Thermo Fisher Scientific, Ref# 4368814). Sample cDNA was diluted 1:10 before input for real-time Q-PCR reaction. Expression of each gene was analyzed with three technical replicates using “PowerSybr Green PCR Master Mix” (Thermo Fisher Scientific, Ref# 4367659) on an Applied Biosystems VIIA7 Q-PCR machine equipped with a 384 well reaction block, and QuantStudio Real-Time PCR Software v1.3. Fold change from gene expression analysis was calculated according to the ΔΔCT method. Primers are listed in the **Supplement Table 6**.

### 5.12 Cell Isolation, Culture, Transfection, PE luciferase assay

Neonatal rat cardiomyocytes (NRCM) were isolated from ∼3-day-old pups from Sprague-Dawley rats (Charles River, Stock 001) following standard protocols using the Worthington Neonatal Cardiomyocyte Isolation System (Worthington, # LK003300) followed by a two-step Percoll density gradient, and cultured in Cardiomyocyte growth media (DMEM/F12 media supplemented with 5% horse serum, 10 mM HEPES, 1% ITS, 15 μg/mL Gentamycin, and 100 μM BRDU).

NRCM were plated at 250K cells per well in 48 well plates, in cardiomyocyte growth media. The following day, NRCMs were co-transfected using Lipofectamine 2000 (0.5% v/v, final) with psiCHECK2 plasmid DNA (20 ng / well) or Txlnb plasmid DNA (200 ng/ well) diluted in Optimem media at a final volume of 100 μL per well. The transfected cells were incubated for four hours and then 150 μL of growth media was added per well. dd

The next day, the media was removed and replaced with serum-free Cardiomyocyte media. After at least 4 hours, phenylephrine was added (100 μM, final concentration in serum-free media) and cells were incubated for ∼ 48 hours, with media exchange after 24 hours. After 48 hours, media was removed and NRCM were lysed/scraped in 100 μl per well of passive lysis buffer, (Promega #E1941, diluted 1:5 with water). Dual-Glo Luciferase assays were performed to measure Firefly and Renilla luciferase activity (Promega #E2940). Lysates (10 μl per well) were loaded into white, opaque 96 well plates in duplicate, and analyzed on a GloMAX 96 well reader with dual injectors. Following a dual injector protocol, where 45 μl of Firefly buffer and 45 μl of Renilla buffer were sequentially added with a 2-second delay and 3-second read time per addition. Raw luciferase data for Firefly and Renilla was analyzed separately, and normalized to the untreated control group. PE-induced luciferase was calculated as PE treated normalized to control treated, and finally, normalized Firefly and Renilla values were averaged together.

### 5.13 Statistical Methods

Data were analyzed for statistical significance using various analyses in Microsoft Excel, GraphPad Prism v10.1.2, and R v3.6.1. Generally, individual replicates are plotted with mean +/- SEM, and statistical significance is directly stated or indicated by convention (* p<0.05, ** p<0.01, *** p<0.001, **** p<0.0001, ns = not significant). Descriptions of specific tests are included in accompanying figure legends. Pairwise comparisons were made using two-tailed T-tests. Comparisons among more than three groups used one-way ANOVA with Sidak posthoc test comparing selected groups, (each comparison is shown on graphs). Survival was compared using a Kaplan-Meier method.

## 6. Acknowledgements.

Txlnb knockout mice were generated at the University of Iowa Genome Editing Core Facility directed by William Paradee, PhD, and supported in part by grants from the NIH and from the Roy J. and Lucille A. Carver College of Medicine. We wish to thank Norma Sinclair, Rongbin Guan, and Joanne Schwarting for their technical expertise in generating transgenic mice. The transgenic Myh6-Cryab^R120G^ mice were graciously donated by Jeffrey Robbins (University of Cincinnati). Echocardiography was performed with the help of the University of Iowa Cardiovascular Phenotyping Core Directed by Dr. Robert Weiss with technical support from Kathy Zimmerman, Alyssa Bosko, and William Kutschke. Experimental research support was provided by Gabrielle Abouassaly, Nathan Witmer, and Henry Lin.

